# Genome-wide analysis identifies *cis*-acting elements regulating mRNA polyadenylation and translation during vertebrate oocyte maturation

**DOI:** 10.1101/712695

**Authors:** Fei Yang, Wei Wang, Murat Cetinbas, Ruslan I. Sadreyev, Michael D. Blower

## Abstract

Changes in gene expression are required to orchestrate changes in cell state during development. Most cells change patterns of gene expression through transcriptional regulation. In contrast, oocytes are transcriptionally silent and use changes in mRNA poly-A tail length to control protein production. Poly-A tail length is positively correlated with translation activation during early development. However, it is not clear how poly-A tail changes affect mRNA translation at a during vertebrate oocyte maturation. We used Tail-seq and polyribosome analysis to measure poly-A tail and translational changes during oocyte maturation in *Xenopus laevis.* We identified large-scale poly-A and translational changes during oocyte maturation and found that poly-A tail changes precede translation changes. Additionally, we identified a family of U-rich sequence elements that are enriched near the polyadenylation signal of polyadenylated and translationally activated mRNAs. A modest density of U-rich elements was correlated with polyadenylation while a high density of U-rich elements was required to activate translation, showing that polyadenylation and translation activation can be uncoupled. Collectively, our data show that changes in mRNA polyadenylation are a key mechanism regulating protein expression during vertebrate oocyte maturation and that these changes are controlled by a spatial code of *cis-*acting sequence elements. Our results provide insight into mechanisms of translational control in oocytes and identify novel proteins important for the completion of meiosis.

## Introduction

Accurate regulation of gene expression is critical for development and cell-fate specification. In most cell types changes in gene expression are controlled at the level of transcription. However, oocytes are transcriptionally silent and changes in gene expression are controlled by posttranscriptional mechanisms [1]. Oocytes complete premeiotic DNA replication and recombination then arrest in prophase of meiosis I [2]. Upon receiving a hormonal signal, oocytes exit the prophase arrest and enter the meiotic divisions. In vertebrates oocytes undergo nuclear envelope breakdown (also called Germinal Vesicle BreakDown, GVBD), segregate homologous chromosomes at meiosis I, extrude a polar body, and arrest at metaphase of meiosis II until fertilization. Completion of these dramatic morphological events requires the coordinated synthesis of many different proteins. Aneuploidy resulting from inaccurate chromosome segregation during the meiotic divisions is a leading cause of birth defects in humans [3], therefore understanding gene expression changes during meiotic maturation is critical to understand the cause of aneuploidy.

Gene expression during oocyte maturation is controlled by cytoplasmic polyadenylation of stored mRNAs [1]. During transcription mRNAs receive a long poly-A tail in the nucleus that facilitates mRNA export. In oocytes the long poly-A tails of most mRNAs are removed in the cytoplasm which triggers ‘masking’ or translational silencing of these mRNAs [4]. As oocytes mature phosphorylation cascades unmask stored mRNAs leading to rapid polyadenylation and polyadenylation-dependent translational activation [5]. Cytoplasmic polyadenylation is facilitated by a *cis-*acting sequence element termed the Cytoplasmic Polyadenylation Element (CPE, UUUUAU) which is bound by the CPEB protein [6]. CPEB and CPEs are important for polyadenylation and polyadenylation-dependent translation of the mos and cyclin B mRNAs during oocyte maturation in *Xenopus laevis [7].* Importantly, cytoplasmic polyadenylation is critical for oocyte maturation as blocking this process completely blocks progress into meiosis [8].

Most of our current knowledge of the mechanisms of cytoplasmic polyadenylation comes from detailed studies of single transcripts. Until recently it has not been possible to measure the length of poly-A tails at a genome-wide scale. In the past 5 years important technical advances have provided the first genome-scale views of changes in transcript polyadenylation [9, 10]. Work from the Bartel and Kim groups have adapted Illumina sequencing to measure poly-A tail length in several experimental systems [9–13]. This work has shown that poly-A tail lengths are carefully regulated during early development in *Drosophila, Xenopus* and zebrafish. In addition, recent work in *Drosophila* has shown that poly-A tail length regulation is important for controlling translation during *Drosophila* oogenesis [11, 14]. Interestingly, poly-A tail length is well-correlated with translational activity during oogenesis and early development, but becomes uncoupled in somatic cells and at the midblastula transition [9]. These recent studies have provided important genome-wide insight into regulation of poly-A tail length during early development and revealed that poly-A tail regulation is a major mechanism controlling gene expression. However, these studies have not determined how poly-A tail lengths are regulated during oocyte maturation in a vertebrate system.

In this work we used multiple genome-wide approaches to measure poly-A tail length and mRNA translation in *X. laevis* oocytes during oocyte maturation. Our results show that poly-A tail length regulation is a major mechanism controlling gene expression in vertebrate oocyte maturation. Additionally, our work provides insight into *cis-*acting sequence elements that control polyadenylation and translation. Our genome-scale approach provides insight into the mechanisms of gene expression control in oocytes and into new proteins important for completion of the meiotic divisions.

## Poly-A tail measurement during oocyte maturation

In order to measure changes in poly-A tail length during *Xenopus laevis* oocyte maturation we implemented the Tail-Seq method in our lab [10]. We synthesized spike-in sequences with defined poly-A lengths (0, 8, 16, 32, 64, 90, 118nt), converted these sequences into an Illumina sequencing library, and sequenced the library. We initially tested the ability of the Illumina base-calling software to correctly infer the length of poly-A stretches in each sample. Consistent with previous results we found that the Illumina base calling software performed poorly on poly-A sequences longer than 32nts (Supplemental Fig. 1A) [10]. We then created a custom algorithm to measure poly-A length based on the detection of the boundary between poly-A sequence and the rest of the transcript. In brief, as opposed to the Illumina fasq sequence files, we used raw fractions of fluorescent intensity corresponding to each base (A,T,G,C) within a 5-nt sliding window around every position of the sequencing read, and detected the shift of these frequencies between adjacent sequence positions (see Methods for details). Our custom poly-A calling algorithm produced much more accurate poly-A length determinations on the synthetic spike-in library (Supplemental Fig. 1B).

To measure genome-wide changes in poly-A tail length we harvested immature oocytes from *X. laevis* females and induced oocyte maturation by the addition of progesterone. We measured meiotic progression by both GVBD and cyclinB:cdk kinase activity. Consistent with previous studies [1], oocytes progressed through meiosis in a stereotypical pattern with two waves of elevated cyclinB:cdk activity representing metaphase of meiosis I and metaphase of meiosis II (Supplemental Fig. 1C). We collected five time points during meiotic progression from two different frogs: untreated (noPG), 90 minutes after progesterone addition (PG90), meiosis I (MI), interkinesis (Int.), and meiosis II (MII). Oocytes from each stage were collected in parallel for analysis of both poly-A tail length and protein translation by sucrose density gradient. We used a modified PAT method [15] to append a 3’ linker to the end of all adenylated mRNAs and convert these mRNAs into a sequencing library that includes intact poly-A tails. We then sequenced libraries from five time points from biological replicates using paired-end sequencing: a 250nt read that sequenced across the poly-A tail and a 50nt read to determine the transcript of origin. We analyzed genes with at least 50 reads in all five time points in both biological replicates (5690 genes, Supplemental Table 1). We found that biological replicates exhibited similar behavior using principal component analysis (PCA) (Fig. 1A), and that the major difference was between samples with inactive (noPG, PG90) and active cyclinB:cdk (MI, Int., MII). We found smaller, reproducible differences between samples as oocytes progressed through meiosis on PC2. Analysis of the average poly-A tail length of transcripts revealed large-scale changes at different stages of meiosis (Fig. 1B). Untreated oocytes exhibited a peak poly-A tail length of ~40nt that was not changed 90 minutes after progesterone addition. However, as oocytes entered MI we observed a large-scale polyadenylation of most mRNAs followed by global deadenylation in Int. and MII (Fig. 1B), consistent with the observation that histone B4 and setd8a mRNAs are polyadenylated then deadenylated during oocyte maturation [16, 17]. Interestingly, a substantial number of mRNAs retained long poly-A tails throughout meiosis. To determine if our TAIL-Seq measurements accurately reflect known patterns of mRNA polyadenylation, we analyzed the poly-A tail lengths of several mRNAs previously shown to be polyadenylated during meiosis (Fig. 1C)[7, 18]. We found that all previously characterized mRNAs exhibited dramatic poly-A tail increases during meiosis. Additionally, we used the PAT assay to measure poly-A tail changes in mRNAs with increased, decreased, and unchanged poly-A tails (Fig. 1F, and Supplemental Fig. 1D). In all tested mRNAs our sequencing results correctly predicted the direction of poly-A tail length change. Taken together these results demonstrate that Tail-Seq accurately captures changes in transcript poly-A tail length during *X. laevis* oocyte meiosis.

**Figure 1.**
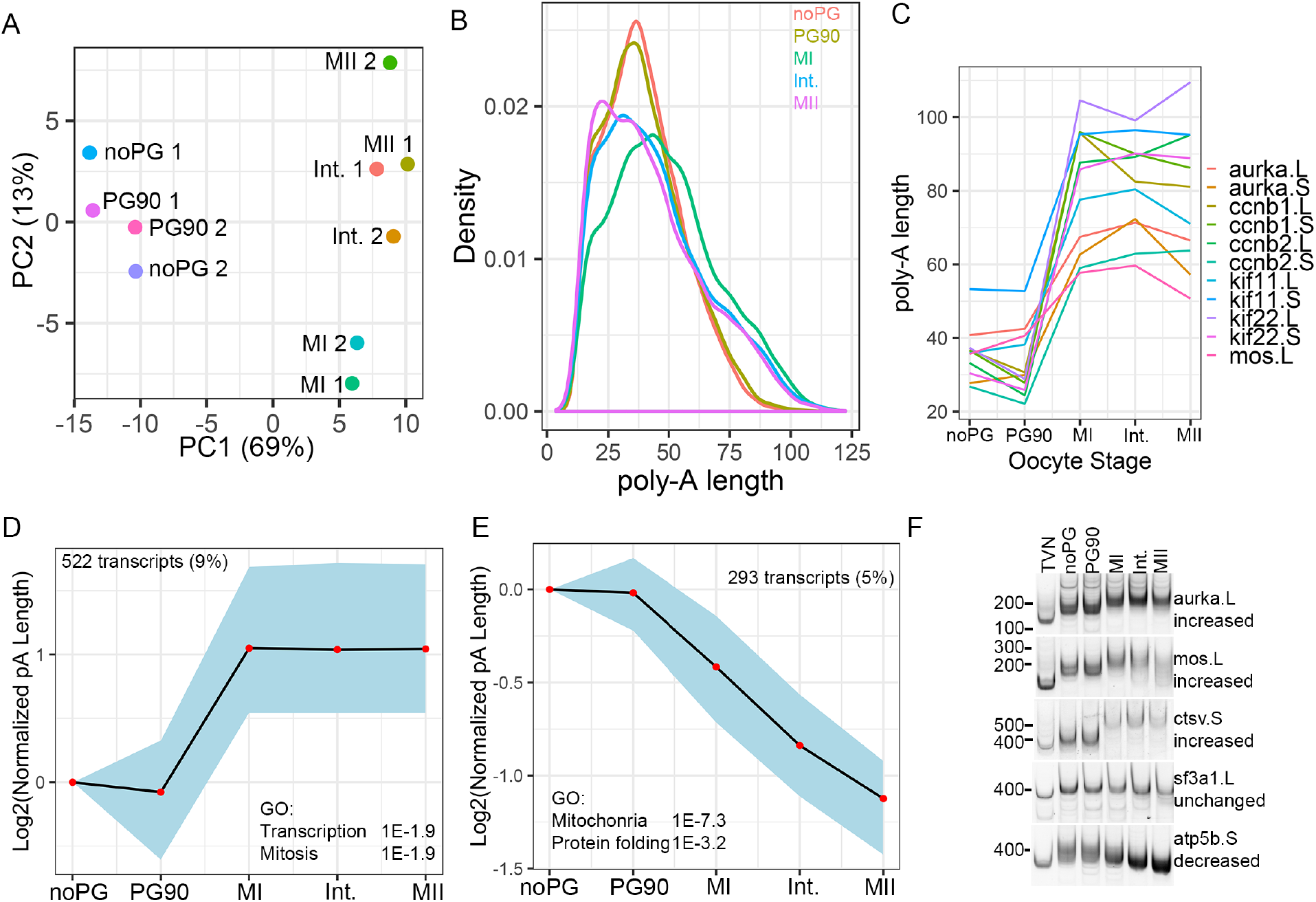
Measurement of poly-A tail changes during oocyte maturation. A. TAIL-seq measurements from two biological replicates were analyzed using Principal Components Analysis. PCA plot shows the relationship between different time points and different biological replicates. B. Histogram showing measured poly-A tail lengths at five time points spanning meiotic maturation. Curves are the average of two replicates. C. Average measured poly-A tail lengths for mRNAs previously reported to be polyadenylated during oocyte maturation. In *X. laevis* many genes have paralogs which are indicated by a .L and .S designation. D. STEM cluster of transcripts exhibiting increased polyadenylation during oocyte maturation. Enriched GO terms are indicated on the plot. E. STEM cluster of deadenylated transcripts and enriched GO terms. F. PAT assay was used to analyze poly-A tail length of the indicated transcripts during oocyte maturation from additional biological replicates (see also Supplemental Fig. 1). Poly-A tail predictions from our TAIL-Seq data are indicated underneath the gene name.

To determine if groups of transcripts exhibited coordinated polyadenylation behavior we used STEM software [19] to cluster transcripts. We identified two major clusters of transcripts that exhibited increased and decreased adenylation during meiosis (Fig. 1DE). 522 transcripts (~9% of measured) increased poly-A tail length at least two-fold during the course of meiosis. Gene Ontology analysis of these transcripts using DAVID software [20] demonstrated that transcripts coding for transcription and mitosis related proteins were enriched in this set. Additionally, 293 transcripts (~5% of measured) exhibited at least two-fold decrease in poly-A tail length. GO analysis demonstrated that transcripts encoding mitochondrial and protein folding proteins were enriched in this cluster (Fig. 1E). We observed that a subset of deadenylated transcripts were degraded during oocyte maturation but that transcript deadenylation was not well correlated with poly-A tail length (Supplemental Figure 2), which is surprising because mRNA deadenylation is one of the first steps of mRNA decay in somatic cells [21]. Our GO analysis is consistent with a study that used Click chemistry to study large-scale changes in poly-A tail length in *Xenopus* oocytes [17]. Collectively, our Tail-Seq results demonstrate that oocytes execute a coordinated program of transcript poly-A tail regulation during oocyte maturation.

**Figure 2.**
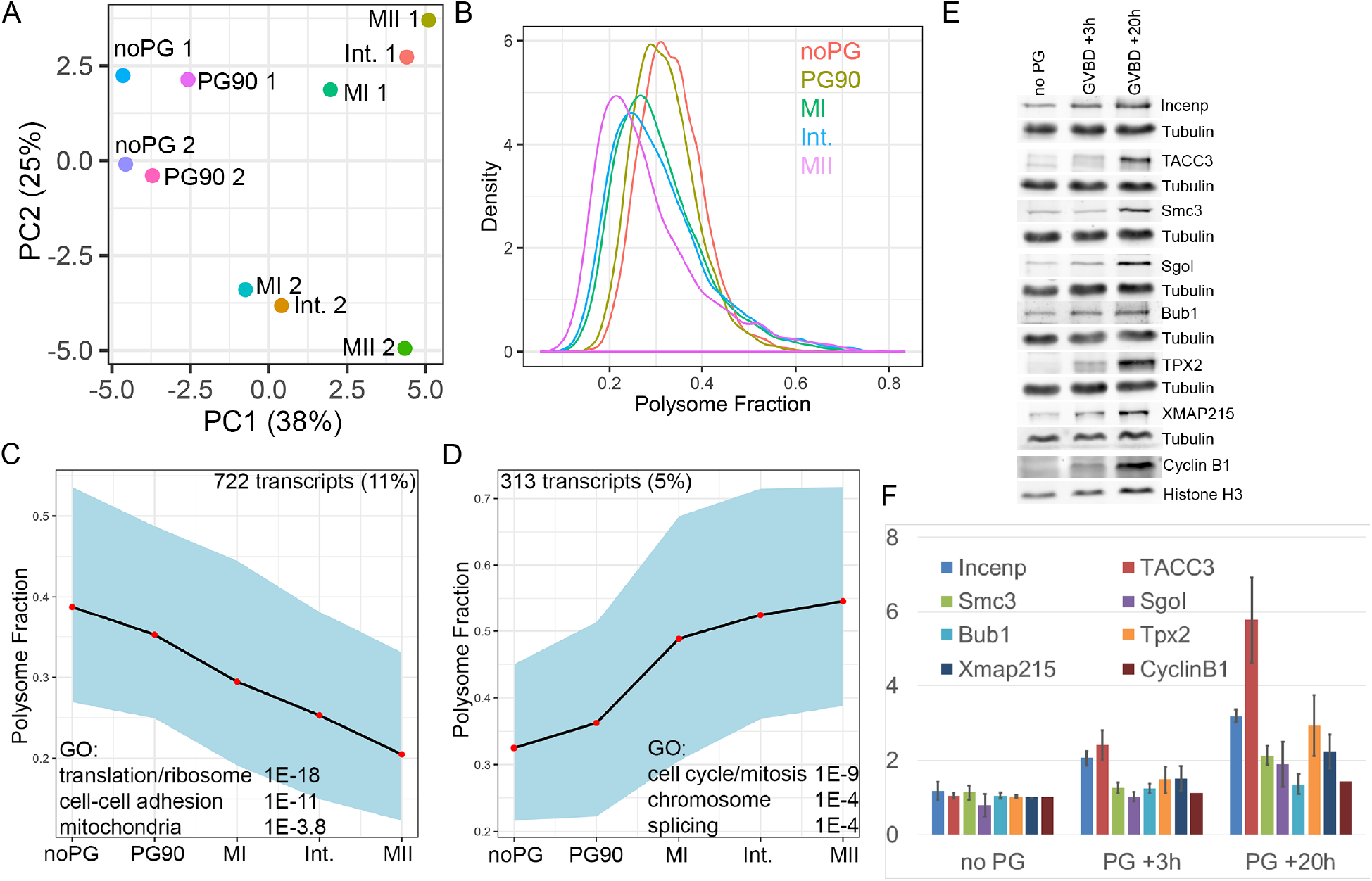
Translational changes during oocyte maturation. A. Extracts from oocytes at different stages of maturation were separated on sucrose gradients and the fraction of mRNA associated with polysomes was calculated. PCA analysis and plot shows the relationship between different oocyte stages and biological replicates. B. Histogram plot showing the fraction of mRNA associated with polysomes at different stages of oocyte maturation. C. STEM cluster of translationally repressed mRNAs and associated GO terms. D. STEM cluster of translationally activated mRNAs and associated GO terms. E. Western blots of candidate proteins predicted to be translationally activated during oocyte maturation. F. Quantitation of Western blots from two biological replicates.

## mRNA translation analysis during oocyte maturation

In *Drosophila* oocytes and several early embryonic systems mRNA poly-A tail length is well-correlated with changes in mRNA translation [9, 11, 14]. Additionally, pioneering work in *X. laevis* oocytes showed that cytoplasmic polyadenylation is necessary to initiate translation of the mos and cyclin B1 mRNAs [7, 16, 22]. To globally determine how changes in mRNA polyadenylation relate to mRNA translation during vertebrate oocyte maturation, we measured mRNA association with polysomes from the same samples as we used to perform Tail-Seq. To determine the efficiency of mRNA translation we separated oocyte extracts on sucrose density gradients and isolated fractions corresponding to: mRNP/40/60S, monosomes, and polysomes (Supplemental Fig. 3). We prepared Illumina libraries from each fraction for each stage of oocyte maturation and calculated the percentage of each mRNA present in the polysome gradient fraction after normalization. We measured the polysome association of 6497 mRNAs using this approach (Supplemental Table 2). PCA analysis demonstrated that different stages of oocyte maturation were reproducibly well-separated by PC1, while PC2 primarily reflected differences between biological replicates (Fig. 2A). Analysis of the average polysome fraction of all transcripts revealed interesting general features of mRNA translation during oocyte maturation (Fig. 2B). We observed a progressive loss of transcripts from the polysome fraction as oocytes progressed through meiosis, consistent with a general repression of translation during M phase [23–25]. Interestingly, we also observed a long tail of transcripts that exhibited increased polysome association during oocyte maturation (Fig. 2B). To verify that changes in mRNA polysome association accurately reflect changes in protein production we analyzed eight candidate proteins that showed increased polysome association by quantitative western blot from additional biological replicates. We found that all eight proteins exhibited significant increases in protein levels during oocyte maturation (Fig. 2EF). In contrast, we also analyzed the protein levels of several proteins that showed decreased mRNA polysome association and were unable to detect a decrease in protein level for any proteins tested (data not shown). This suggests that these proteins may be stable during oogenesis, and decreases in mRNA association with polysomes do not lead to rapid changes in protein levels unless it is coupled with active proteolysis of the preexisting protein.

**Figure 3.**
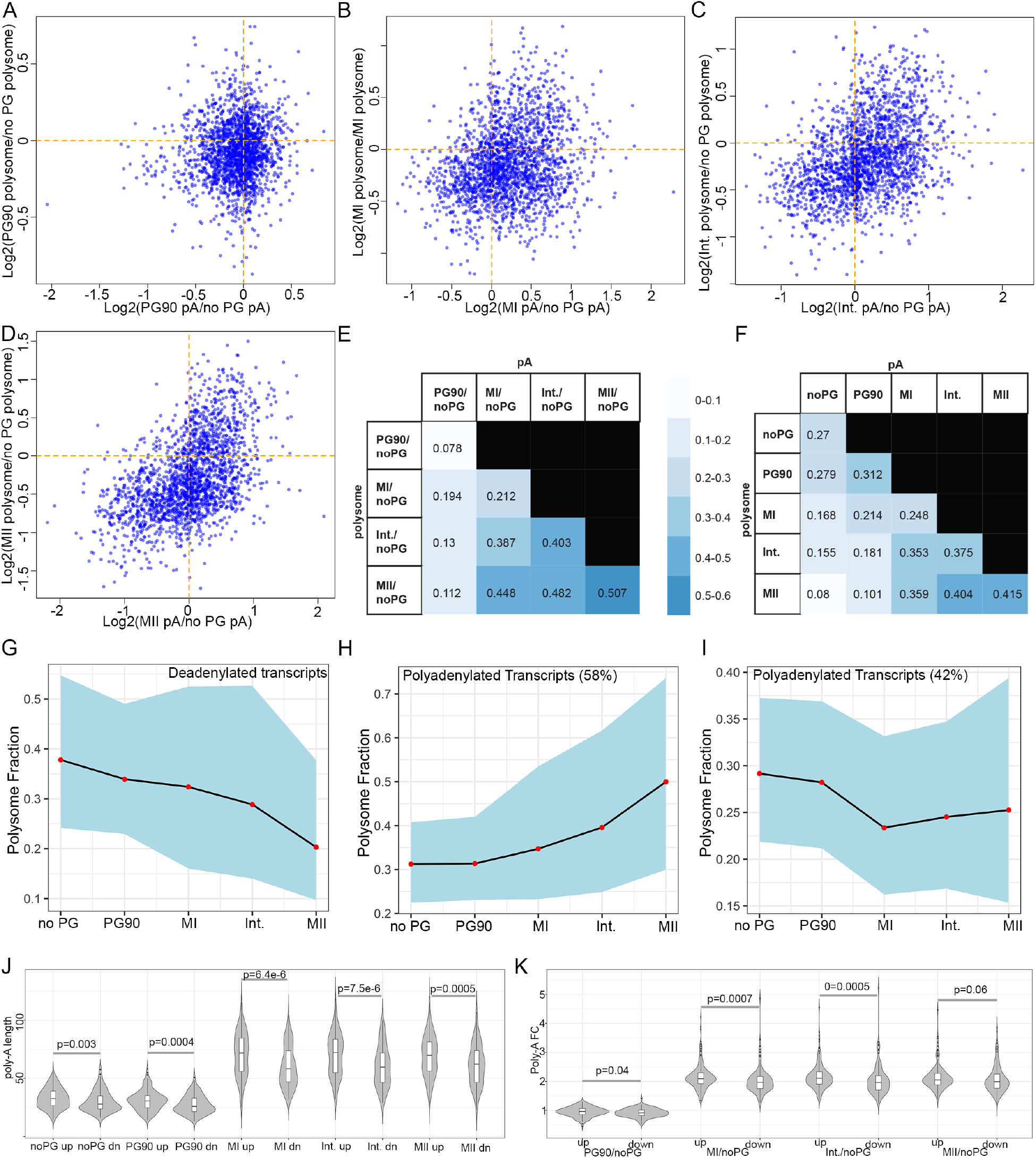
Correlation between transcript polyadenylation and translation. A-D. Scatterplots of the change in polyadenylation compared to the change of the fraction of mRNA present on polysomes. E. Spearman correlation coefficients between changes in polyadenylation and changes in mRNA fraction on polysomes. F. Spearman correlation coefficients of raw poly-A tail length and raw polysome percentage measurements. G. The translation behavior of deadenylated transcripts (from Fig. 1E) was examined using STEM software. All deadenylated transcripts exhibited translational repression. H-I. The translation behavior of adenylated transcripts (From Fig. 1D) was examined using STEM software. Adenylated transcripts were either translationally activated (H) or repressed (I). J. Violin plots of the raw poly-A tail lengths of polyadenylated transcripts that were translationally activated (up) or repressed (down, dn). K. Violin plots of the fold change in poly-A tail length of adenylated transcripts that were translationally activated (up) or repressed (down). Indicated p-values are the result of a Wilcox test.

To determine if oocytes change the translated fraction of specific classes of transcripts we performed STEM clustering of mRNAs based on translation behavior. We identified two major (and 4 minor) clusters of mRNA translation behavior (Fig. 2CD, Supplemental Fig. 3). The first cluster contains 722 transcripts (~11% of measured) and exhibits a dramatic decrease in fraction of mRNA present on polysomes (Fig. 2C). GO analysis demonstrates that ribosomal, cell-cell adhesion, and mitochondrial protein transcripts are highly enriched, similar to the GO categories enriched in deadenylated and degraded (Supplemental Fig. 2) mRNAs. Cluster 2 contains 313 transcripts (~5% of measured) and exhibits a significant increase in mRNA fraction present on polysomes (Fig. 2D). GO analysis demonstrates that cell cycle/mitosis and splicing proteins are highly enriched in this cluster. Taken together these results show that oocytes regulate mRNA translation as they mature and that specific functional classes of transcripts are up-or down-regulated.

### Complex relationship between polyadenylation and translation

To understand the relationship between transcript polyadenylation and translation we combined our Tail-Seq and polysome data. We analyzed 2093 transcripts where we had reliable data for both poly-A tail length and polysome fraction in both biological replicates (Supplemental Table 4). We analyzed the correlation between changes in poly-A tail length and changes in the fraction of mRNA on polysomes across oocyte maturation (Fig. 3A-E). At early stages of meiosis changes in poly-A tail length and translation were very weakly correlated, but the correlation between the two factors improved as oocytes progressed through maturation (Fig. 3E). In general, increases in poly-A tail length were positively correlated with an increased fraction of mRNA on polysomes and decreased poly-A tail length positively correlated with a decreased mRNA polysome fraction. Interestingly we observed a clear temporal offset in correlation between poly-A changes and translation changes (Fig. 3E), which is consistent with studies of mos and cyclin B1 during *X. laevis* oocyte maturation [16]. This is especially well-illustrated when comparing changes in poly-A tail length in MI to changes in translation at MI, Int. and MII. The poly-A tail changes are nearly always better correlated with polysome changes at a later developmental stage, suggesting that poly-A tail changes precede translational changes. Additionally, we examined the correlation between raw poly-A tail length and fraction of mRNA present on polysomes (Fig. 3F). We observed that poly-A length was also positively correlated with polysome fraction, suggesting that both raw poly-A tail length and changes in poly-A tail length contribute to changes in mRNA translation.

To determine how changes in polyadenylation affect translation for a defined set of transcripts we examined the translational behavior of clusters of mRNAs exhibiting increased or decreased polyadenylation (Fig. 1DE) during oogenesis. We used STEM software to cluster transcripts with decreased poly-A tail length based on translational behavior (Fig. 3G). We found that all deadenylated transcripts had reduced mRNA levels on polysomes (Fig. 3G). In contrast, when we clustered polyadenylated transcripts based on translational behavior we found two distinct groups of translational behavior (Fig. 3HI). First, 58% of transcripts showed both increased poly-A tail lengths and increased translation, consistent with a positive correlation between longer poly-A tails and increased translation. Surprisingly, we also identified a second cluster (42% of transcripts) that showed unchanged or decreased polysome association despite having increased poly-A tails. Interestingly, polyadenylated mRNAs with unchanged translation exhibited significantly shorter poly-A tails than translationally activated mRNAs, suggesting that there may be a poly-A tail length threshold necessary to strongly activated translation (Fig. 3JH). Taken together, our results indicate that mRNA deadenylation always leads to decreased polysome association of mRNAs. In contrast, mRNA polyadenylation can lead to both unchanged and increased translation, suggesting that additional mRNA features may control translation in addition to mRNA poly-A tail length.

### Genome-wide identification of polyadenylation sites and prediction of 3’UTR sequences

In order to understand if sequence elements located in 3’UTRs control the polyadenylation and translation behavior of mRNAs, we undertook an analysis of mRNA 3’UTRs. Many genes have multiple possible polyadenylation sites (PASs) whose use can differ based on developmental context [26–29]. In order to accurately annotate PASs in *X. laevis* oocytes we performed PAS-Seq [30] in Stage VI oocytes. After read filtering and peak calling we identified 42,859 PASs supported by >= 4 independent reads. 41,982 PASs (98%, Supplemental Table 5) mapped to within 5kb downstream of annotated genes (Fig. 4D), demonstrating the utility of PAS-Seq to correctly identify the 3’ end of transcripts and improve mRNA annotations. The majority of PASs mapped to 3’UTRs (Fig. 4A). PAS sites mapped to 13,216 genes with the majority of transcripts having a single PAS site (Fig. 4B, E). However, many transcripts clearly contained multiple PAS sites (Fig. 4F) that are used in differing ratios. For all subsequent analysis we calculated PAS usage by counting the number of reads supporting each PAS and only considering the 3’UTRs corresponding to PASs with the greatest number of reads (11,055). In many cases this lead to a clear single dominant PAS (Fig. 4D), while the usage of other PASs was more equally balanced (Fig. 4F).

**Figure 4.**
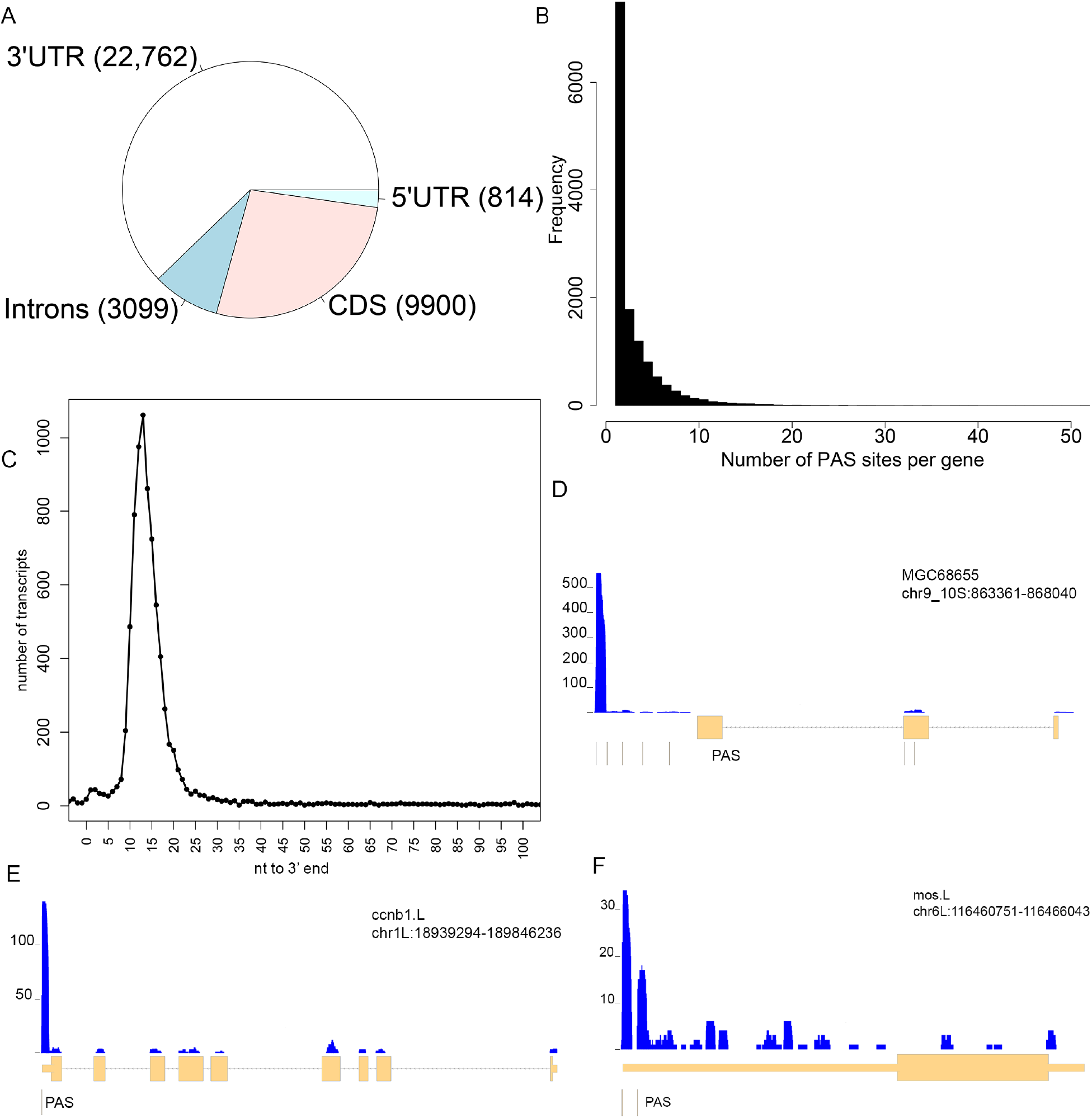
PAS-Seq analysis of the *X. laevis* oocyte transcriptome. A. PAS-Seq was used to identify polyadenylation sites in *X. laevis* oocytes. Pie chart shows that fraction of PASs mapping to each type of transcript feature. B. Histogram of the number of PASs per gene. C. Histogram of the average number of nucleotides from the 3’ most base of the polyadenylation signal (AAUAAA, AUUAAA) to the measured end of the transcript. D-E. Genome browser views of PAS-seq reads compared to three genes. PAS calls are listed below each gene as thin bars.

To explore the general characteristics of PASs we searched for the presence of a canonical polyadenylation signal (poly (A) signal) (AAUAAA or AUUAAA) and found that 83% contained at least one poly (A) signal. Additionally, we determined the distance from the most 3’ polyadenylation signal to the measured 3’ end of the transcript. As observed with human RNAs [31] we found that most RNAs contained a polyadenylation signal < 20nt from the 3’ end of the RNA (Fig. 4C). Collectively, PAS-Seq allowed us to experimentally identify 3’ ends of transcripts in *X. laevis* oocytes and to accurately predict 3’ UTR sequences for analysis.

### 3’UTR length and primary sequence motifs control transcript polyadenylation and translation

To determine what transcript features influence polyadenylation and translation in oocytes we analyzed the length and composition of motifs in experimentally determined 3’UTR sequences. A recent study found that senescent mouse cells use distal poly(A) (pA) sites, leading to a global lengthening of 3’ UTRs and reduced gene expression [32]. To study if the length of 3’UTRs can modulate translation of polyadenylated mRNAs, we examined 3’UTR length in clusters of transcripts with differential association with polysomes. We found that transcripts with increased polyadenylation but slow or unchanged translation exhibited significantly longer 3’UTR sequences compared to mRNAs that were adenylated and translationally activated (Fig. 5A). Longer 3’ UTRs are likely to contain increased numbers of regulatory elements that could decrease translation. Interestingly, deadenylated and translationally repressed transcripts did not exhibit longer 3’UTR sequences. Additionally, translationally activated and repressed transcripts (Fig. 2CD, Fig. 5A) did not exhibit significant differences in 3’ UTR length suggesting that multiple mechanisms control mRNA translation.

**Figure 5.**
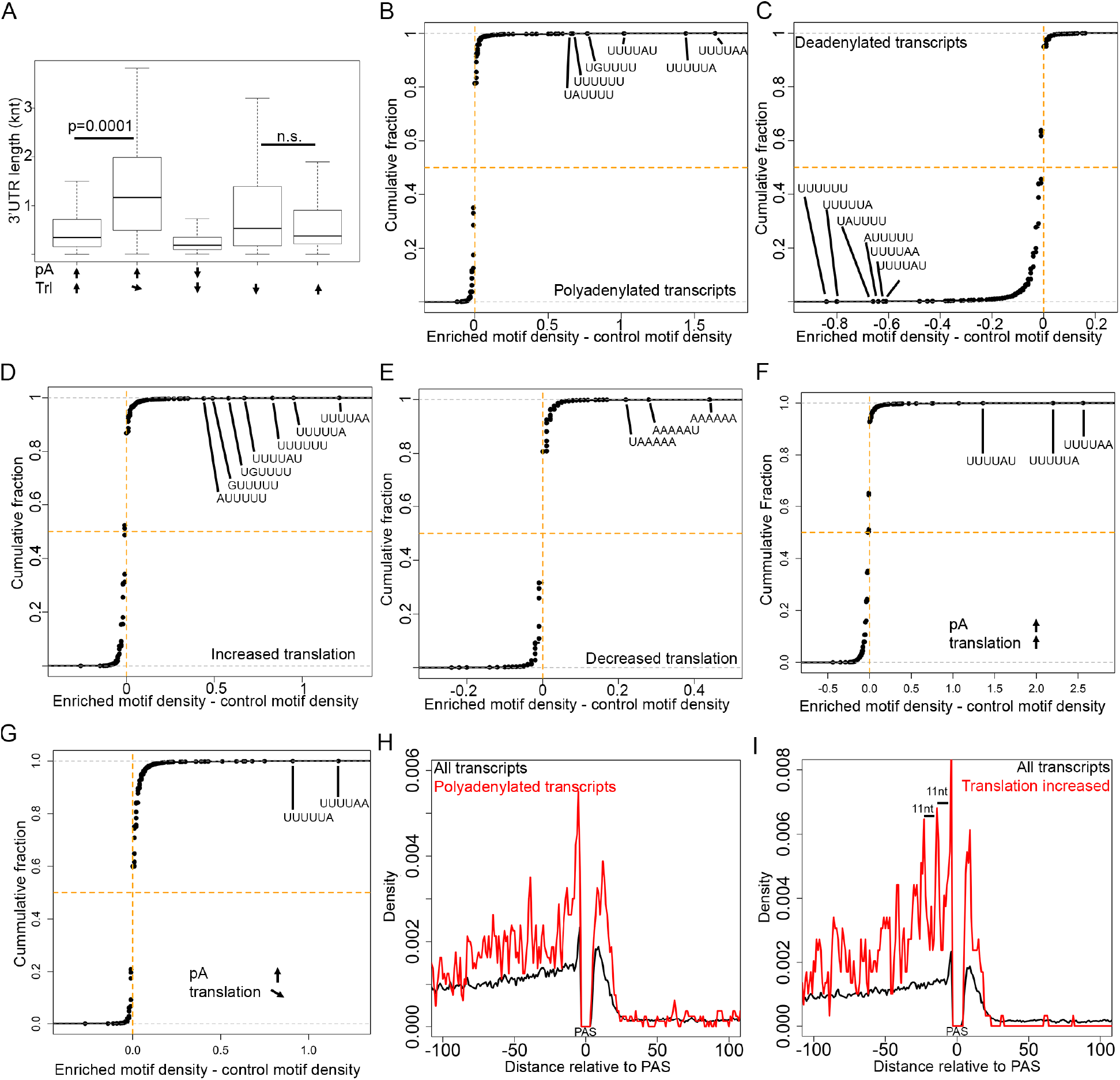
Primary sequence motifs are enriched in polyadenylated and translationally activated transcripts. A. 3’UTRs lengths were measured from clusters of mRNAs exhibiting various polyadenylation and translation behavior. P-values are the result of a Wilcox test. B-E. The density of all hexamers was calculated in the 3’UTRs of various sets of transcripts (polyadenylated (B), deadenylated (C), translationally activated (D), translationally repressed (E)) and in all transcripts. Cumulative distribution plots show the enrichment of various hexamers in different subsets of transcripts compared to all transcripts. F-G. Hexamer enrichment was calculated in the 3’UTRs of polyadenylated transcripts that were translationally activated (F) or translationally repressed (G). Translationally activated transcripts exhibit a higher density of U-rich sequence elements. H. The positions of the top 13 U-rich elements (from B) were compared to the 3’-most polyadenylation signal for all transcripts (black line) and adenylated transcripts. I. The relative density of U-rich sequences compared to the 3’-most polyadenylation signal for translationally activated transcripts.

To determine if specific sequence elements are enriched in clusters of transcripts with coordinated behavior we calculated the enrichment of all hexamers [33] (either as density or as a Z-score) in clusters of transcripts compared to all transcripts. We first compared the groups of transcripts that were polyadenylated or deadenylated during oocyte maturation. A large family of U-rich hexamers were enriched in polyadenylated transcripts (Fig. 5B, Supplemental Table 6). The U-rich motifs are similar to and include the CPE element (UUUUAA or UUUAAU) which is well-documented to promote transcript polyadenylation, suggesting that these motifs are likely binding sites for CPEB1 [1, 34]. In contrast, deadenylated transcripts did not contain any enriched hexamers (Fig. 5C, Supplemental Table 7), but exhibited a significant depletion of the family of U-rich hexamers. Collectively these data suggest a model for how *cis-*acting sequence elements control transcript polyadenylation. A high density (or specific location, see below) of U-rich sequence elements promotes transcript polyadenylation while an absence of U-rich elements promotes transcript deadenylation. This data suggests that transcript deadenylation is the default behavior during oocyte maturation and that deadenylation is counteracted by the presence of U-rich sequences in the 3’UTR.

To determine if *cis-*acting sequence elements also influence the translation behavior of mRNAs we calculated hexamer enrichment in translationally up- and down-regulated mRNAs. Similar to polyadenylated transcripts a large family of U-rich hexamers were also enriched in translationally activated mRNAs (Fig. 5D, Supplemental Table 9), with the CPE element being the most enriched element. In contrast, only a few A-rich motifs were very modestly enriched in translationally repressed mRNAs (Fig. 5E, Supplemental Table 8). To determine if the density of U-rich elements affects the translation behavior of polyadenylated mRNAs we compared the density of U-rich hexamers in mRNAs that are polyadenylated and translationally activated (Fig. 3H, Fig. 5F) or translationally repressed (Fig. 3I, Fig. 5G). Interestingly we found that the density of U-rich elements is much higher in translationally activated mRNAs compared to translationally repressed mRNAs. Collectively, these results suggest that a high density of U-rich sequence elements promote both polyadenylation and polyadenylation-dependent translational activation, and a low density of U-rich elements promote polyadenylation but not translation activation, while A-rich elements may function to recruit translational repressors. These results are consistent with the strong correlation between transcript polyadenylation and translation (Fig. 3) and the known role of the CPE element in promoting polyadenylation and translation. Additionally, the identification of simple sequence motifs is consistent with a recent analysis of the preferred binding motifs of a large number of RNA-binding proteins [34].

Extensive analysis of the role of CPE elements in the regulation of polyadenylation and translation has demonstrated that the location of the CPE relative to the polyadenylation signal and to other CPE elements has a strong influence on CPE functionality [22, 35]. To determine if U-rich sequence elements are located at specific positions in groups of mRNAs with coordinated behaviors we analyzed the location of U-rich hexamers relative to the poly-A signal. U-rich sequences were modestly enriched immediately before and after the polyadenylation signal in all transcripts (Fig. 5HI black line), but showed no enrichment at other positions. In contrast, polyadenylated transcripts exhibited strong peaks of U-rich elements before and after the polyadenylation signal as well as multiple secondary peaks 5’ of the poly-A signal (Fig. 5H). Translationally activated transcripts also exhibited a strong enrichment of U-rich elements before and after the polyadenylation signal (Fig. 5I). Additionally, translationally activated mRNAs exhibited several phased peaks 5’ of the polyadenylation signal that are separated by ~11nt. The appearance of phased U-rich elements fits well with the observation that CPE sites separated by 10-12nt lead to translational repression of cyclin mRNAs before oocyte maturation, and that the distance between the CPE and poly-A signal regulates polyadenylation-dependent translational activation [22]. Collectively, our data suggest that U-rich sequence elements are enriched near the polyadenylation signal of mRNAs that are polyadenylated and translated during oocyte maturation. Additionally, U-rich element density and spacing likely contribute to polyadenylation and translational activation (Fig. 6).

**Figure 6.**
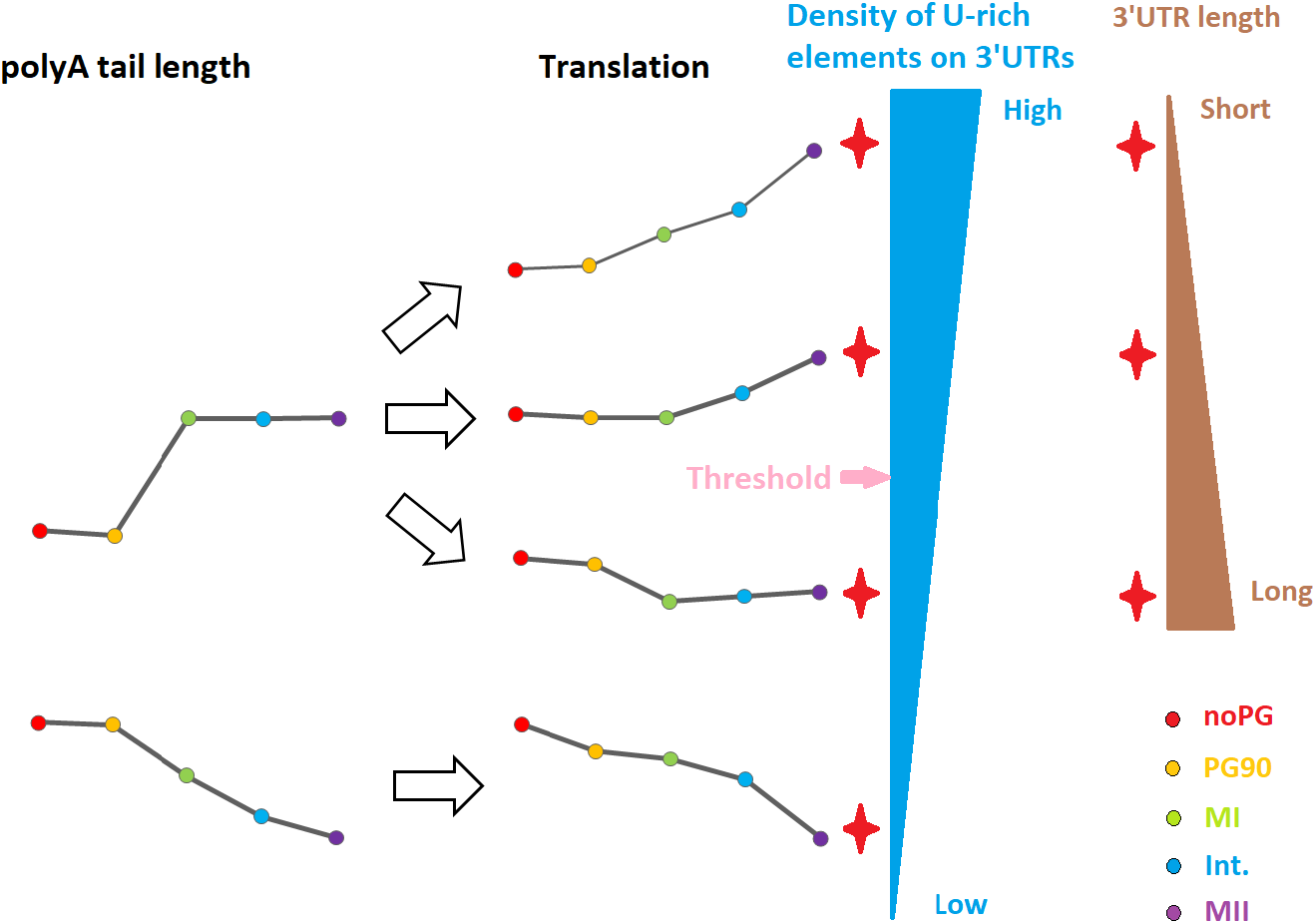
Model for control of polyadenylation and translation by U-rich sequence elements and 3’UTR length.

## Discussion

We used a combination of genome-wide approaches to provide a high temporal resolution view of transcript polyadenylation and translation during vertebrate oocyte maturation. We found large-scale changes in both poly-A tail lengths and translation, suggesting that oocytes execute a precise posttranscriptional gene expression program to complete the meiotic divisions and prepare for early development following fertilization. Our work identified *cis-*acting sequence elements whose density may set a threshold for activation of polyadenylation-dependent translation and is likely a rich source of novel proteins that facilitate oocyte maturation (Fig. 6).

### Large-scale changes in poly-A tail length during oocyte maturation

Several recent genome-wide studies demonstrated that large-scale changes in transcript polyadenylation occur during oogenesis and early development [9–11, 13, 14]. In particular, two recent studies found large changes to poly-A tail length and a strong correlation between changes to poly-A tail length and changes in translation in *Drosophila* oocytes [11, 14]. However, it is not possible to experimentally control the progress of oocytes through maturation in *Drosophila* and it was not clear how changes in poly-A tail length are related to the morphological events of oocyte maturation. In addition, *Drosophila* oocytes arrest in metaphase of meiosis I prior to fertilization while vertebrate oocytes arrest at metaphase of meiosis II, so it is not clear if *Drosophila* is an accurate model of vertebrate oocyte maturation. We confirmed that a large fraction of the transcriptome changes poly-A tail length during oocyte maturation. We found that most of the transcriptome is polyadenylated as oocytes undergo GVBD and that the majority of transcripts are then deadenylated as oocytes progress through the meiotic divisions. Global transcript polyadenylation is similar to the global polyadenylation observed when comparing mature and immature *Drosophila* oocytes [11]. Interestingly, we found that a subset of transcripts that code for proteins associated with mitosis retain long poly-A tails through oocyte maturation. As a result of the experimental tractability of *X. laevis* oocytes we were able to obtain a high temporal resolution view of poly-A and translation changes during oocyte maturation. We found that changes in poly-A tail length precede changes in translation. Collectively, our work adds to the growing body of evidence that global poly-A tail length changes are a highly-conserved feature of oocyte maturation and are used to control gene expression in transcriptionally silent cells.

Pioneering work in *X. laevis* demonstrated that *cis-*acting sequence elements, such as the CPE, control cytoplasmic polyadenylation in *Xenopus* oocytes [7, 36]. In depth analysis of the mos and cyclin B1 mRNAs demonstrated that cytoplasmic polyadenylation is strongly influenced by the number and positions of the CPEs element relative to the polyadenylation signal [22, 35]. However, it has not been possible to determine if there is a genome-wide correlation of the CPE element with cytoplasmic polyadenylation or to determine how the CPE positions globally influence cytoplasmic polyadenylation. To address this issue we used PAS-Seq to identify the polyadenylation sites of transcripts in immature oocytes and experimentally defined 3’UTR to search for motifs enriched in polyadenylated and deadenylated transcripts. Interestingly, we found a large family of U-rich sequence motifs that were enriched both before and after the polyadenylation signal in polyadenylated transcripts. Additionally, we could detect secondary enrichment peaks for U-rich hexamers, consistent with previous work suggesting that clusters of precisely spaced CPE elements promote efficient cytoplasmic polyadenylation [22]. The fact that we identified many different U-rich motifs, including the CPE, suggests that CPEB proteins likely can bind to a range of U-rich sequences, which is consistent with recent work [34]. Surprisingly, we found that no sequence motifs were enriched in deadenylated transcripts, but rather that deadenylated transcripts were characterized by an absence of U-rich motifs. Taken together these results suggests that transcript deadenylation is the default behavior as oocytes mature and that deadenylation is counteracted by the presence of CPE-like sequences near the polyadenylation signal. Deadenylation as a default behavior has previously been proposed to explain the behavior of ribosomal protein mRNAs during oocyte maturation on the basis of careful examination of individual transcripts [37]. Our work confirms this hypothesis and extends this principle to a genome-wide scale.

### Translation changes during oocyte maturation

To determine if poly-A tail length and translation are correlated during vertebrate oocyte maturation we measured the fraction of each mRNA associated with polyribosomes. We found that translation was globally repressed as oocytes progressed through the meiotic divisions, consistent with longstanding observations from somatic cells that protein translation is globally repressed during M phase [23–25]. However, we identified a large group of transcripts that increased translation during oocyte maturation including a group of transcripts coding for proteins involved in completion of meiosis. A large number of transcripts coding for proteins involved in mRNA splicing are also translationally activated, which is surprising because oocytes are transcriptionally silent [38]. It is tempting to speculate that splicing components are upregulated in preparation for the onset of zygotic transcription. Alternatively, it is possible that splicing components are upregulated to promote the cytoplasmic splicing of transcripts with retained introns.

Extensive work studying the CPE element has shown properly spaced CPE elements serve to translationally repress mRNAs prior to oocyte maturation and promote “early” (prophase), “late” (metaphase I), or “late late” translational activation following oocyte activation [22]. We found that a family of U-rich elements were enriched immediately before and after the polyadenylation signal in translationally activated mRNAs, which is predicted to activate “early” and “late” translational activation [22]. Additionally, we found strong, regularly spaced peaks of U-rich sequence elements at a spacing of ~11nt, which is strikingly similar to the optimal spacing of CPE elements found in cyclin mRNAs to promote translational repression prior to maturation and translational activation during maturation. Our results suggest that the same family of U-rich sequence elements is the major global mechanism that controls both mRNA polyadenylation and translation during oocyte maturation. Additionally, we detected a weak enrichment of A-rich elements in translationally repressed mRNA, suggesting that these elements may recruit translational repressors to some mRNAs.

### Coordinated regulation of polyadenylation and translation during oocyte maturation

During oogenesis and early development poly-A tail length and mRNA translation are well correlated in *Drosophila, Xenopus*, and zebrafish. Our results are consistent with previous studies in that we find that poly-A tail length and changes to poly-A tail length are well correlated with changes in mRNA translation during vertebrate oocyte maturation. Interestingly, we find that poly-A tail length is uncorrelated with translation in prophase-arrested oocytes and becomes more correlated as oocytes progress through meiosis. We found that changes in poly-A tail length were better correlated with changes in translation than raw poly-A tail length. Importantly, our high temporal resolution dataset allowed us to observe a temporal offset in changes in poly-A tail length and changes in translation. This suggests that lengthening or shortening poly-A tails causes changes in translation initiation.

Examination of the translational behavior of polyadenylated and deadenylated groups of transcripts revealed important information about the relationship between the two processes. First, deadenylated transcripts were always translationally repressed, suggesting that transcript deadenylation and translational repression are the default regulation as oocytes progress through meiosis. Second, adenylated transcripts exhibit complex translational behavior. Consistent with work on the cyclin B and mos mRNAs a majority of polyadenylated transcripts are translationally activated. However, a substantial fraction of polyadenylated transcripts show no change in translation or are translationally repressed. The major difference between translationally activated and unchanged transcripts is the density of U-rich elements. This novel result supports that hypothesis that the density and spacing of U-rich elements is the critical determinant for controlling translation. Previous work demonstrated that polyadenylation is always correlated with translational activation, but our work shows that a high density of U-rich elements is necessary to activated translation on polyadenylated transcripts. Low density of non-optimally spaced U-rich elements can promote polyadenylation, but the correct spacing is critical to promote translational activation.

In summary, we have found that large scale changes in protein expression occur during vertebrate oocyte maturation and that these changes are strongly coupled to changes in poly-A tail lengths. Our work confirms work in other systems and suggests that cytoplasmic polyadenylation is the major mechanism regulating protein expression in transcriptionally silent cells. In addition, our work has identified a large group of transcripts that are translationally activated during oocyte maturation that could be a rich source of novel proteins important for the completion of oocyte meiosis.

## Experimental Procedures

### Xenopus oocytes preparation and maturation in vitro

Female frogs [39] were primed with 50 U of chorionic gonadotropin human (hCG) 5-14 days before the oocyte collection. On the day before oocyte collection, fresh filtered 1×Modified Barth’s Solution (1×MBS) was prepared, and stored at 4°C. Frogs were anesthetized and euthanized by an overdose of MS222 [40] followed by decapitation and ovaries are collected and soaked in 1×MBS. Then ovaries were washed, cut into small pieces (1-2cm^2^), and defolliculated with Liberase TL (0.025mg/ml) with gently shaking at 23°C overnight. The reaction was stopped by adding 1×MBS, and oocytes were rinsed with 1×MBS for several times and stage VI oocytes were collected using an 800μm filter. Stage VI oocytes were matured by addition of 3μM progesterone in 1×MBS at 21°C or 23°C. Samples were collected at a series of time points (No progesterone, progesterone 90min, GVBD 0min, GVBD 30min, GVBD 60min, GVBD 90min, GVBD 120min, GVBD 150min, GVBD 180min) during oogenesis, then flash frozen in liquid nitrogen and stored at −80°C for: H1 kinase assay, TAIL-seq, analysis of polysomes, and PAT assay. For each time point, 5 oocytes were used for H1 kinase assay, TAIL-seq and PAT assay respectively. The oocytes used for TAIL- seq and analysis of polysomes were from the same two frogs, but biological replicates were utilized for PAT assay. For each type of experiment, two biological repeats were performed using two different frogs.

### H1 Kinase assay

Five oocytes were homogenized in 100μL of EB (80 mM β-glycerophosphate, 20mM EGTA, 15mM MgCl_2_) + leupeptin, pepstatin, cymostatin and phosphatase inhibitors (type I and type II, Sigma-Aldrich). Lysates were centrifuged at 20K × g for 15 minutes at 4°C and clear extract was collected and stored at −80°C. Kinase assays were performed by incubating 5μL of lysate with Histone H1 and γ-^32^P ATP in EB for 15 minutes. Reactions were stopped by the addition of SDS loading buffer and separated using a 15% SDS-PAGE. Gels were exposed to phosphor-screens for ~3 hours and scanned using a GE Typhoon imager.

### TAIL-seq

Based on H1 kinase results, oocytes at five time points (No progesterone, progesterone 90min, GVBD 0min, GVBD 90min, GVBD 150min for repeat 1 or 180min for repeat 2) during oogenesis were picked for RNA extraction and RNA-seq library construction. First, oocytes were crushed by pellet pestles in 40ul of XB buffer (100 mM KCl, 0.1 mM CaCl_2_, 1 mM MgCl_2_, 50 mM sucrose, 10 mM HEPES at pH 7.7), then 500ul Trizol was added to extract total RNAs from whole oocytes. Before adding chloroform, yolk and debris were removed by centrifuge (12000g, 5min at 4°C). The total RNAs in aqueous phase were extracted again using Direct-zol RNA miniPrep Kit. TAIL-seq libraries were constructed based on Harrison et al.’s method [15] with some modifications: Briefly, the 3’end of RNA was extended by Klenow polymerase exo- at 37°C for 1h, and secondary structures were disrupted by adding 1×digest buffer and heating at 80°C for 10min. The extended RNAs were fragmented with 0.1U/ul RNase T1 at 37°C for 15min, and the reaction was stopped by phenol/chloroform extraction. After binding to streptavidin beads, the cDNAs were 5’-phosphorylated with T4 PNK, ligated with 5’ splinted linker, reverse transcribed using PAT-seq end-extend primer, eluted from streptavidin beads by 30ul of 0.13% SDS with heating at 100°C for 5min and then sitting on ice for 2min. The eluted cDNAs libraries were size selected twice by equal volume of AMPure XP beads (Beckman Coulter, Inc.), and amplified for 14 cycles by NEBNext Ultra II Q5 Master Mix (NEB), NEBNext Universal PCR primer for Illumina (5’-AAT GAT ACG GCG ACC ACC GAG ATC TAC ACT CTT TCC CTA CAC GAC GCT CTT CCG ATC-s-T-3’) and NEBNext Index primer for Illumina (NEB, index# 9-13 for five libraries in each repeat). Then the amplified cDNAs libraries were purified again by equal volume of AMPure XP beads (Beckman Coulter, Inc.), and 1ul of eluted libraries were taken for analysis by Bioanalyzer. The size of PAT-seq libraries were 150-1000bp, with average size from 389bp to 423bp. For each biological replicate, we mixed the five libraries at a same molar percentage (19.6%), together with 4 spikes of 0, 8, 16 and 32 As (with index #8-5) at a same molar percentage (0.1%) and 4 spikes of 64, 80, 90 and 118 As (with index #4-1) at a same molar percentage (0.4%). The libraries mixtures were run paired-end Illumina Hiseq with 2 full lanes in rapid mode: 50 cycles reading from the 5’sq end (P5), and 250 cycles from the 3’sq end (P7).

### TAIL-Seq Sequencing Analysis

Using intensity CIF files and cluster location CLOCS files produced by the Illumina HiSeq instrument in addition to the standard FASTQ files, we extracted flowcell coordinates and intensity of each passing sequencing cluster in the four acquisition channels (A, C, G and T). For each sequencing position of read 2 in each library fragment, we calculated the normalized intensities for each of the four channels by normalizing these four values to the total of 1. We then used these normalized intensities for the positions of the sequencing read to estimate the length of poly-T tail within each read. Similar to the previous observation [10], we observed that in the control samples with the known length of poly-T tail, the end of the poly-T tail coincides with a rapid reduction of signal intensity for the T channel and a rapid increase in the signal intensities for non-T channels. To detect these changes along the read length, we developed and optimized a relatively simple and robust computational approach based on the analysis of signal intensities within a sliding window. For each read 2 of the sequencing library, we used a sliding window of 8 bp with the step of 1 bp to calculate the mean normalized intensity for each base type within this window and applied the following rule to determine the end of poly-T tail:

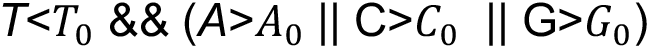

where *A*,*T*,*G*,*C* are average signal intensities of each base in the window and *A*_0_, *T*_0_, *G*_0_, *C*_0_ are the cutoff values of the corresponding intensities. We called the end of the polyT tail when this rule was satisfied at least three times among 5 consecutive positions of the sliding window. The cutoff values *A*_0_, *T*_0_, *G*_0_, *C*_0_ were optimized on the control samples with known lengths of poly-A tails. We used the following cutoffs: *A*_0_ = 11, *T*_0_ = 45, *G*_0_ = 11, *C*_0_ = 30 Read 1 on the opposite end of the library fragment was mapped to the *Xenopus laevis* transcriptome XL_9.1_v1.8.3.2 using STAR aligner [41]. The average poly-A tail length for each transcript was estimated by calculating the mean length of poly-T tails among all library fragments with read 1 mapped to a given transcript.

### Sucrose density gradient and polysome library preparation

Sucrose gradients were prepared in 50mM Tris pH 7.5, 250mM KCl, 25mM MgCl_2_. Gradients were poured in steps and frozen in liquid nitrogen between steps. Gradients were stored at −80°C and thawed overnight at 4°C before use. Oocytes (~100) from each stage were incubated in polysome buffer (20mM Tris pH 7.5, 0.5% NP-40, 300mM KCl, 2mM MgCl_2_ with 0.1mg/ml cycloheximide for 5 minutes at room temperature. Oocytes were then allowed to settle and buffer was removed. Oocytes were crushed with a pestle and centrifuged at 17K × g for 15 minutes at 4°C. Cytoplasmic layer was removed for use in sucrose gradients. 25μL of cytoplasmic extract was diluted with 250μl of PB containing cycloheximide and loaded onto a thawed sucrose gradient. Gradients were centrifuged at 39000 rpm using a SW41 rotor for 2 hours at 4°C. Tubes were pierced using a Brandel tube piercer and fractionated by upwards displacement using a GE Akta FPLC. 500μL fractions spanning the entire gradient were collected and stored at −80°C.

Total RNA was purified from each fraction using Trizol. RNA samples from mRNP, monosome, and polyribosomes was pooled and digested with RQ1 DNase. DNase-treated RNA was cleaned up using the Zymo Clean & Concentrate kit. 100ng of total RNA from human RPE-1 cells was added to each sample and poly-A RNA was purified using Exiqon LNA dT kit. Poly-A RNA was then used as input for the NEBNext Ultra Directional Library Prep Kit for Illumina according the manufacturer’s instructions. Libraries were PAGE purified (we selected 250-600bp range) and sequenced on an Illumina HiSeq.

### Polysome sequence alignment and analysis

Reads were collapsed into unique sequences using a custom Perl script [42]. Unique reads were aligned to Xenbase mRNA sequences using Bowtie2 or to the *Xenopus laevis* 9.2 genome release using Tophat2 [43]. In addition, unique reads were aligned to human Refseq RNAs using Bowtie2. Reads were counted against mRNAs using a custom Perl script [42] or the cuffdiff function of cufflinks. FPKM values for all human genes detected at a FPKM of >10 were used to normalize FPKM values between libraries. For each sucrose gradient we used normalized FPKM values to calculate the percentage of mRNA present in the polysome fraction of the gradient. Polysome percentages were used to analyze changes in mRNA translation during oocyte maturation. We analyzed mRNAs that could be detected at a FPKM >1 in all samples from both biological replicates. All sequences from polysome gradients and PAS-Seq were deposited in GEO with accession number GSE134537.

### RNA degradation analysis

To measure transcript stability during oocyte maturation we used reads generated from our sucrose gradient experiments. We merged fastq files from each gradient fraction for each time point to create a ‘total RNA’ file for each timepoint. We then used Tophat2 and Cufflinks to align and count these reads against the *X. laevis* genome. We searched for reproducibly up- and down-regulated RNAs using STEM software.

### PAT assay and TVN PCR

Based on H1 kinase results, oocytes from 5 time points (No progesterone, progesterone 90min, GVBD 0min, GVBD 120min for repeat1 or 150min for repeat 2, GVBD 180min) during oogenesis were selected and total RNA was purified using Trizol. Using USB Poly (A) Tail-length Assay Kit (Affymetrix, USB), 0.5ug total RNAs were used for each 10ul of G/I tailing reaction, then the reaction was stopped and 7.5ul were reverse transcribed, and 2.5ul were used as a negative control with no reverse transcription. NEBNext Ultra II Q5 Master Mix (NEB) was used to amplify diluted poly(G/I) tailed cDNAs with PAT Universal Reverse Primer (5’-GGTAATACGACTCACTATAGCGAGACCCCCCCCCCTT-3’) and forward gene specific primer. TVN PCR was used as a control of PAT assay: 2.5ug total RNAs per 20ul reaction were reversed transcribed at 50C for 1h, using TVN PCR reverse primer (5’-CAAGCAGAAGACGGCATACGATTTTTTTTTTTTTTTTTTVN-3’) and SuperScript III Reverse Transcriptase (Invitrogen), and then the reaction was stopped by heating at 70°C for 15min. To reduce non-specific amplification, touch-down PCR program (98°C, 30s; [98°C, 10s, 72°C, 75s]_7 cycles, −1°C each cycle_; [98°C, 10s; 65°C, 75s]_23cycels_, 65°C, 5min; 12°C,end) were used to amplify cDNAs from PAT assay and TVN PCR. Finally, PCR products from PAT assay and TVN PCR were run on same native TBE gels and stained with SYBR® Gold Nucleic Acid Gel Stain (Invitrogen) for analysis.

### PAS-seq library preparation

Total RNAs extracted from stage VI oocytes (no PG) were used for preparation of PAS-seq libraries from three biological replicates. However, during library sequencing barcode reading failed and we could not separate the three libraries, therefore they were treated as a single replicate. PAS-Seq primers are listed below:

HITS-3 (5’-ACA CTC TTT CCC TAC ACG ACG CTC TTC CGA TCT TTT TTT TTT TTT TTT TTT TVN NN-3’), HITS-5 (5’-CGG TCT CGG CAT TCC TGC TGA ACC GCT CTT CCG ATC TrGrG rG-3’), PA1.0 (5’-AAT GAT ACG GCG ACC ACC GAG ATC TAC ACT CTT TCC CTA CAC GAC GCT CTT CCG ATC TTT TTT CTT TTT TCT TTT TT-3’), PAIndex 1(5’-CAA GCA GAA GAC GGC ATA CGA GAT ATC ACG CGG TCT CGG CAT TCC TGC TGA ACC GCT CTT CCG ATC T-3’), PAIndex2 (5’-CAA GCA GAA GAC GGC ATA CGA GAT CGA TGT CGG TCT CGG CAT TCC TGC TGA ACC GCT CTT CCG ATC T-3’), PAIndex3 (5’-CAA GCA GAA GAC GGC ATA CGA GAT TTA GGC CGG TCT CGG CAT TCC TGC TGA ACC GCT CTT CCG ATC T-3’), Sequencing primer (5’-ACA CTC TTT CCC TAC ACG ACG CTC TTC CGA TCT TTT TTC TTT TTT CTT TTT T-3’).

### PAS-Seq sequence alignment and data analysis

Reads from two Hi-Seq lanes were combined into a single fastq file and collapsed into unique reads. Linker sequences were removed using the fastx toolkit keeping all trimmed reads 30nt or longer. Reads were aligned to *X. laevis* 9.2 genome assembly using Bowtie2. Aligned reads were filtered to retail alignments that contained 2 or more nontemplated As, originating from the poly-A tail. We then used Yodel peak calling software (https://github.com/LancePalmerStJude/YODEL) to convert aligned PAS-Seq reads into peaks. Yodel was used with default settings and required a minimum peak height of 4. All PAS-Seq reads and poly-A site calls are deposited in GEO under accession GSE134537.

### Prediction of 3’UTRs based on PAS-seq

We first predicted 42859 polyA sites located at genome, based on PAS-Seq read peaks called using Yodel software (reads>=4). We used the following criteria: if reads are aligned to the + strand of genome, polyA site should be at the 5’-most of peak aligned region; if reads are aligned to the – strand of genome, polyA site should be at the 3’-most of peak aligned region. Next, we determined that 41982 polyA sites intersected with gene models ‘XENLA_9.2_Xenbase.gff3’, using the following criteria: (1) PolyA site should be located at the same chromosome or scaffold as the gene. (2) The alignment of polyA site to genome should be located at the opposite strand of the gene. (3) The polyA site should be located in the range from start of the gene to 5 kb downstream of the gene. Using gene model ‘XENLA_9.2_Xenbase.gff3’, we found 46582 *X. laevis* gene with stop codons, and picked 43793 genes with unique stop codon to locate polyA sites and predict their corresponding 3’UTRs. For 3’UTR prediction, we chose 22762 polyA sites after the unique stop codon, and only one mRNA with annotated most exons after stop codon for each gene.

3’UTRs are from the first nucleotide after the stop codon to the polyA site on exons. 22762 of 3’UTRs are predicted following the following criteria: (1) For genes on + strand, polyA site is after stop codon with two conditions: (1.1) No exons are between polyA site and stop codon. 3’UTR sequence is all nucleotides between (stop codon, polyA site]. (1.2) Exons are between polyA site and stop codon, also with two conditions: (1.2.1) polyA site is between the two ends of an exon. 3’UTR sequence is all exons nucleotides between (stop codon, polyA site]. (1.2.2) polyA site is between two exons. 3’UTR sequence is all exons nucleotides between (stop codon, polyA site], together with the intron nucleotides between [last valid exon above, polyA site]. (2) For genes on – strand, polyA site is before stop codon with two conditions: (2.1) No exons are between polyA site and stop codon. 3’UTR is reverse complement of all nucleotides between [polyA site, stop codon). (2.2) Exons are between polyA site and stop codon with two conditions. (2.2.1) polyA site is between two ends of an exon. 3’UTR is reverse complement of all exons nucleotides between [polyA site, stop codon). (2.2.2) polyA site is between two exons. 3’UTR is reverse complement of the intron nucleotide between [polyA site, first valid exon above], together with all exons nucleotides between [polyA site, stop codon). ‘[’ and ‘]’ means being included, ‘(’ and ‘)’ means not being included. Finally, we picked 11055 of 3’UTRs with most read numbers for each gene for future analysis.

### Distributions of polyA sites on genes based on PAS-seq

For 41982 polyA sites located on gene model, we obtained 22762 polyA sites after and 13934 polyA sites before the stop codon of mRNAs used for 3’UTRs prediction. We then picked 2789 genes with multiple stop codons, and located 5017 polyA sites on these genes. The remaining 269 polyA sites are found on genes with no stop codon (MT genes, lncRNA genes etc.). For the 13934 polyA sites before the stop codon, we located them on 5’UTRs, CDSs and introns.

Briefly, for each polyA site, we first got their exons and CDSs information of their corresponding mRNAs in ‘XENLA_9.2_Xenbase.gff3’. Then we arranged exons and CDSs in an order from small position to big one, based on their left and right end value. Next, we defined introns and 5’UTRs using following criteria: Introns is all nucleotides between (right end of exon, left end of next exon). For mRNAs on + strand, 5’UTRs is all nucleotides between [left end of the first exon, the left end of the first CDS); for mRNAs on – strand, 5’UTRs is all nucleotides between (right end of last cds, right end of to last exon]. ‘[’ and ‘]’ means being included, ‘(’ and ‘)’ means not being included. Finally, we located 814 polyA sites on 5’UTRs, 9900 on CDSs and 3099 on introns. And there are 121 remaining polyA sites with 113 on mRNAs which are not the ones used for 3’UTRs prediction, and 8 on mRNAs with no stop codon.

### Sequence motif analysis

To analyze enriched motifs we used 3’UTR sequences predicted from our PAS-Seq data. For mRNAs with multiple PAS sites we analyzed the sequence supported by the largest number of PAS reads (11055 sequences). To analyze hexamer composition we calculated that density of all hexamers in various clusters of UTRs (from Figures 1, 2, and 3) and all transcripts. We then calculated hexamer enrichment by subtracting the mean hexamer density in all transcripts from the mean density in a cluster of transcripts. For each hexamer we also calculated a Z-score for motif density and calculated the percentage of 3’UTRs that contained the motif. We repeated this analysis with motif lengths from 4nt to 8nt and observed similar results.

To analyze the relative position of U-rich motifs in 3’UTRs we identified all sequences that contained a match to a canonical polyadenylation sequence (AAUAAA or AUUAAA) and retained the most 3’ polyadenylation location. We then calculated the distance of all matches to the top 13 U-rich motifs to the most 3’ polyadenylation signal for various clusters of RNAs and for all transcripts.

### Data access

All raw and processed sequencing data generated in this study have been submitted to the NCBI Gene Expression Omnibus (GEO; http://www.ncbi.nlm.nih.gov/geo/) under accession number GSE134537.

## Acknowledgements

We thank Ulandt Kim and the Molecular Biology sequencing core for help with optimization of the Tail-seq procedure, Adrian Salic for the gift of Sgo1 antibodies, Judith Sharp and Carlos Perea-Resa for comments on the manuscript. This work was funded by a grant to M.D.B. from NIH (GM108548) and by NIH/NIDDK P30 DK040561 to R.I.S.

---

**Supplemental Figure 1.**
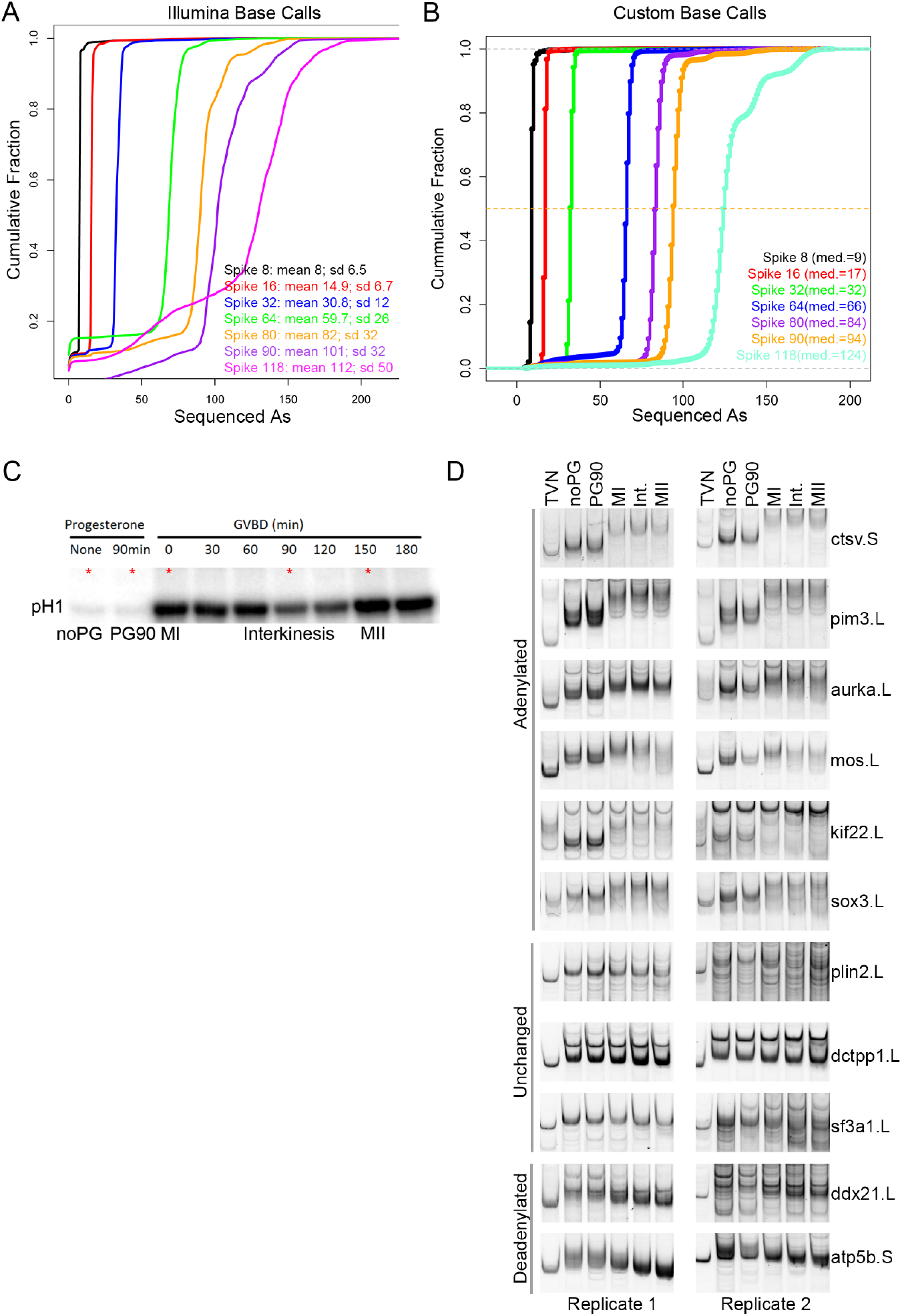
TAIL-Seq validation. A. Synthetic spike sequences containing various lengths of A homopolymers were sequenced using an Illumina miSeq. Cumulative distribution plot shows the poly-A lengths as determined by the Illumina base-calling software. B. Custom software was developed to more accurately determine the length of poly-A sequences from raw Illumina sequencing files. Cumulative distribution plot shows the distribution of poly-A lengths determined by our custom software on the same sequencing data presented in A. C. Histone H1 kinase assays of oocytes as they progress through meiotic maturation. Red Asterix indicates time points that were used for TAIL-seq and polysome analysis. D. PAT assay was used to measure poly-A tail length of mRNAs that were predicted to be polyadenylated, unchanged, or deadenylated by our TAIL-Seq data. Two biological replicates are included for each gene.

**Supplemental Figure 2.**
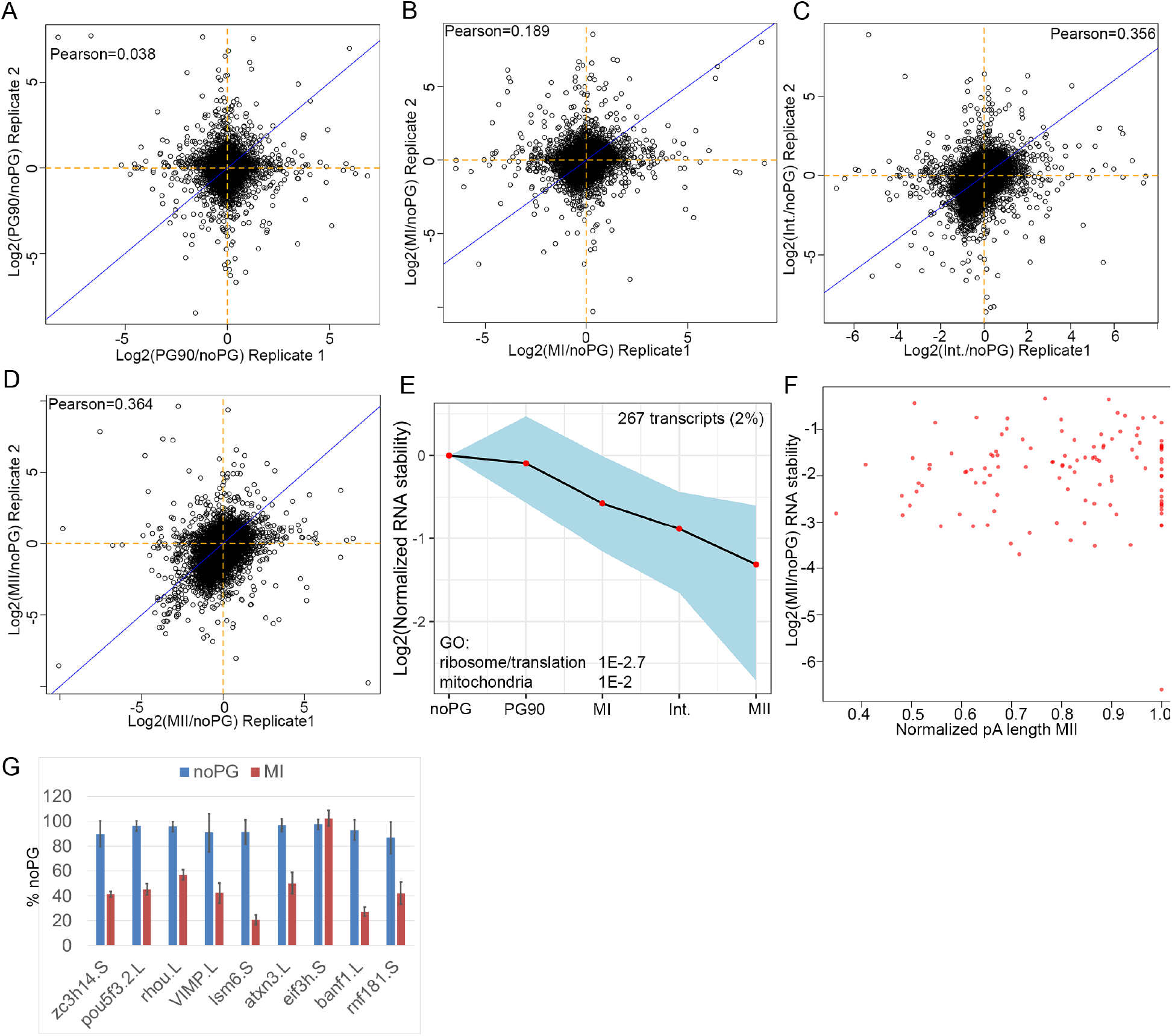
RNA degradation during oocyte maturation. RNA-seq reads from polysome gradients were pooled to create ‘total RNA’ samples for analysis of transcript stability during oocyte maturation. A-D. Scatterplots comparing two biological replicates of changes in RNA abundance during oocyte maturation. E. STEM software was used to identify clusters of transcripts that were up- or down-regulated during oocyte maturation. One cluster of degraded transcripts was identified. GO terms enriched in this set of transcripts are indicated on plot. F. Scatterplot comparing change in RNA stability for degraded transcripts (from E) to normalized measured poly-A tail length. G. Q-RT-PCR was used to measure the stability of 9 transcripts predicted to be degraded from two additional biological replicates. We also tested several transcripts that were predicted to be increased during oocyte maturation and were unable to confirm any changes in transcript levels.

**Supplemental Figure 3.**
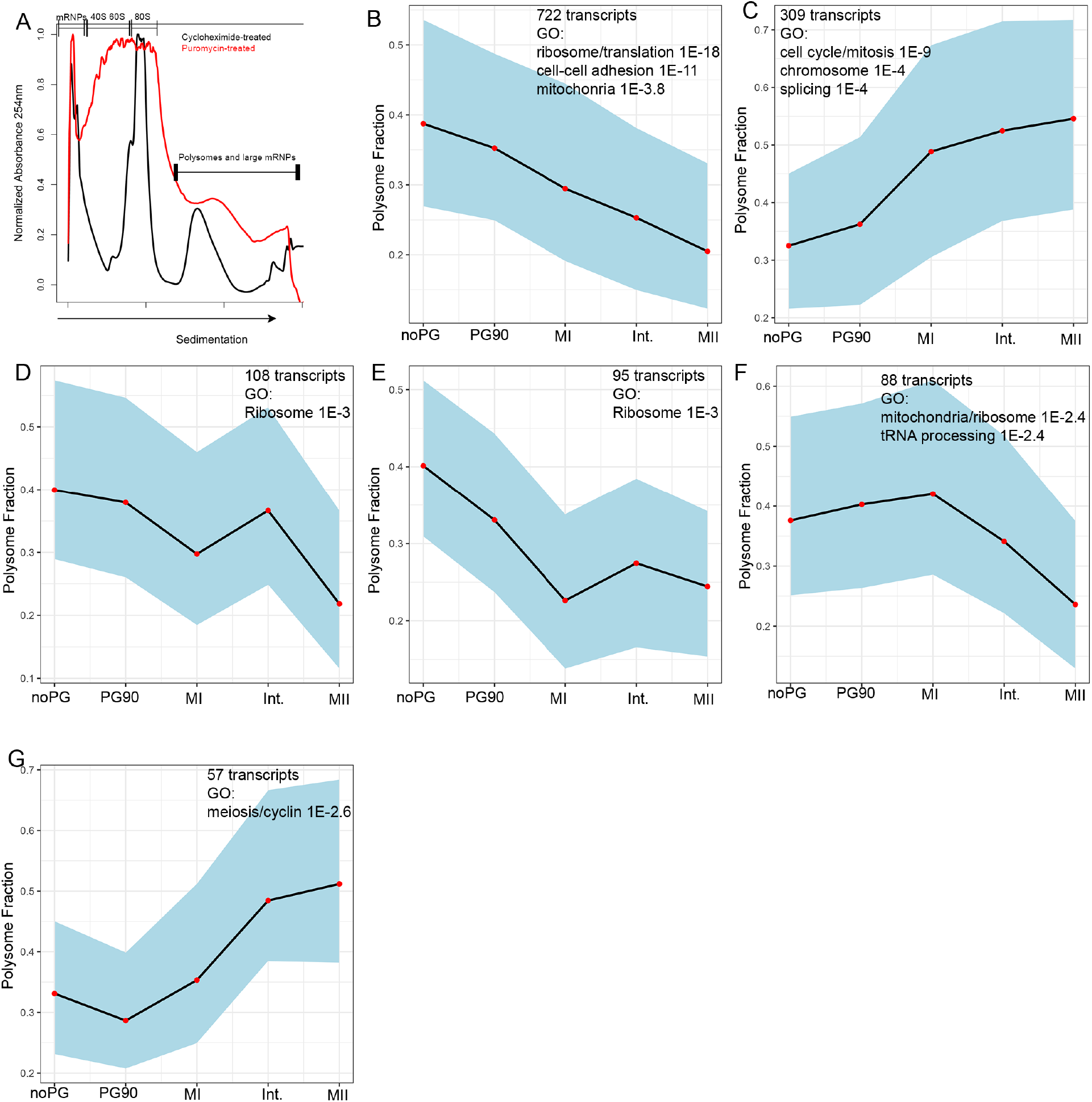
Additional analysis of translation during oocyte maturation. A. Oocyte extract incubated with cycloheximide (black) or puromycin (red) was separated on a sucrose density gradient and UV traces were collected. Different RNA fractions are indicated above the gradient. B-G. Translational changes were clustered using STEM software. Plots indicate the translational behavior of each cluster, the number of genes in each cluster, and the enriched GO terms for each cluster.

**Supplemental Table S1. Poly-A tail measurements for all transcripts with at least 50 reads**.

**Supplemental Table S2. RNA stability measurements.**

**Supplemental Table S3. Polysome fraction of each transcript**.

**Supplemental Table S4. Combined poly-A and polysome percentage measurements.**

**Supplemental Table S5. PAS site calls from PAS-Seq**.

**Supplemental Table S6. Hexamer enrichment from polyadenylated transcripts.**

**Supplemental Table S7. Hexamer enrichment from deadenylated transcripts.**

**Supplemental Table S8. Hexamer enrichment from translationally activated transcripts.**

**Supplemental Table S9. Hexamer enrichment from translationally repressed transcripts**.

